# *MEPSi*: A tool for simulating tomograms of membrane-embedded proteins

**DOI:** 10.1101/2022.07.27.501771

**Authors:** Borja Rodríguez de Francisco, Armel Bezault, Xiao-Ping Xu, Dorit Hanein, Niels Volkmann

**Author notes:** Corresponding author: Niels Volkmann.

## Abstract

The throughput and fidelity of cryogenic cellular electron tomography (cryo-ET) is constantly increasing through advances in cryogenic electron microscope hardware, direct electron detection devices, and powerful image processing algorithms. However, the need for careful optimization of sample preparations and for access to expensive, high-end equipment, make cryo-ET a costly and time-consuming technique. Generally, only after the last step of the cryo-ET workflow, when reconstructed tomograms are available, it becomes clear whether the chosen imaging parameters were suitable for a specific type of sample in order to answer a specific biological question. Tools for *a-priory* assessment of the feasibility of samples to answer biological questions and how to optimize imaging parameters to do so would be a major advantage. Here we describe *MEPSi* (*M*embrane *E*mbedded *P*rotein *Si*mulator), a simulation tool aimed at rapid and convenient evaluation and optimization of cryo-ET data acquisition parameters for studies of transmembrane proteins in their native environment. We demonstrate the utility of *MEPSi* by showing how to detangle the influence of different data collection parameters and different orientations in respect to tilt axis and electron beam for two examples: (1) simulated plasma membranes with embedded single-pass transmembrane αIIbβ3 integrin receptors and (2) simulated virus membranes with embedded SARS-CoV-2 spike proteins.

**HIGHLIGHTS:**

- Tool to simulate tomograms of membrane-embedded proteins
- Detangles influence of data acquisition parameters from sample quality issues
- Rapid evaluation and optimization of cryo-ET data acquisition parameters
- Proof-of-concept provided with integrins and SARS-CoV-2 spike simulations

**GRAPHICAL ABSTRACT:** 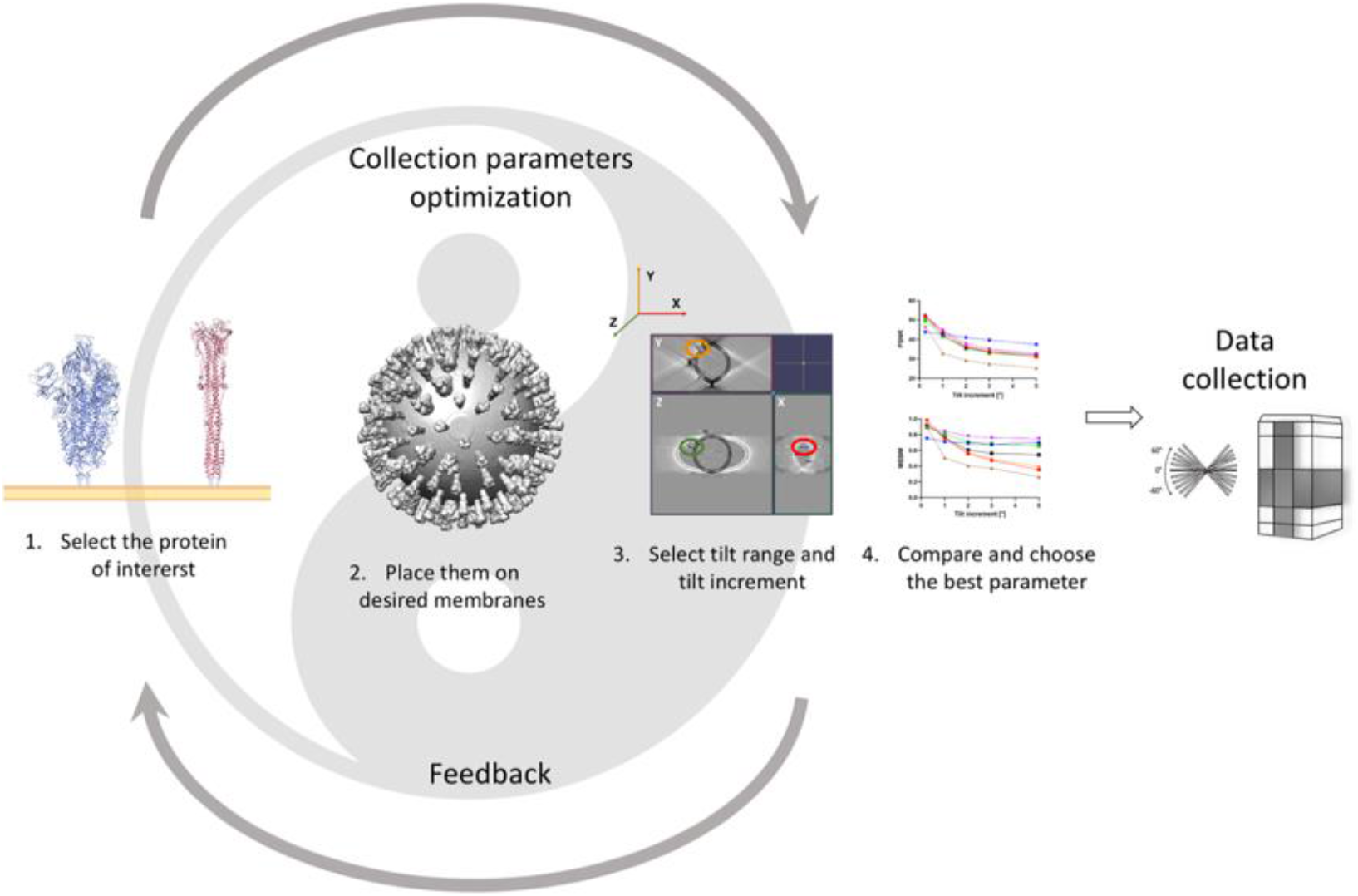

## INTRODUCTION

Since the ‘resolution revolution’ (Kühlbrandt, 2014) cryogenic electron microscopy structure deposition in specialized databases has grown exponentially. In fact, its growth rate is significantly higher than X-ray crystallography or nuclear magnetic resonance spectroscopy (Sorzano and Carazo, 2022). This trend has been made possible because of a growing number of cryo-EM facilities as well as optimized sample preparation and image processing procedures. However, these studies are restricted to purified proteins and protein assemblies. The interest in *in-situ* structural information is increasing because proteins in their native cellular environment may not always behave similar to those *in vitro* purified systems. Cryo-ET is a state-of-the-art technique that, in principle, can provide such data. In combination with averaging techniques, cryo-ET can even enable near-atomic resolution structure determination of specific macromolecular complexes within intact cells (Tegunov et al., 2021). However, sample preparation protocols, correlation with light microscopy data, setup of appropriate data collection parameters, and image reconstruction strategies all need to be carefully optimized and verified to yield tomographic reconstructions or sub-tomogram averages that can answer specific biological questions. This process is time consuming and costly.

One way to improve upon the current process is the introduction of simulation studies to help determine and optimize appropriate parameters. By detangling effects owing to insufficient sample quality and to choices of data collection parameters and image reconstruction strategies for ideal samples, simulations can help bracketing the expectations of what can be achieved with ideal samples prior to investing in imaging and sample optimization. For example, if the simulations indicate that it is not feasible to extract the targeted biological information from ideal samples, there is no point to invest in sample optimization and imaging. Furthermore, as simulations can be cheaper than experiments, a large number of conditions and combinations can be tested and evaluated to maximize sample usage and data collection time. Finally, simulations potentially allow to detangle the effects of parameters that are not experimentally separable (e.g., missing wedge versus tilt increment), allowing a more complete understanding of the influence of the individual parameters.

Simulations have been extensively used in various imaging fields to evaluate image processing and reconstruction algorithms. It was in 1974 when the Shepp-Logan phantom was developed, a tool to simulate the head and brain for computerized tomography (Shepp and Logan, 1974). Since then, the Shepp-Logan phantom has been used as a standard test for evaluating image acquisition and reconstruction algorithms in many different fields. For cellular cryo-ET, early simulations have been primarily used to validate template matching and sub-tomogram averaging approaches (Bartesaghi et al., 2008; Förster et al., 2008) or the effect of typical cryo-ET artifacts on molecular docking (Volkmann, 2002). More recent efforts have focused on improving the accuracy of the image formation model (Himes and Grigorieff, 2021; Parkhurst et al., 2021; Vulović et al., 2013; Rullgård et al., 2011; Hall et al., 2011), which is thought to be especially crucial for cellular tomography, owing to the thickness of the sample, the crowded environment inside cells, and the fact that tilting of large fields of view can lead to major focus differences within the sample. For the most part, underlying models designed to emulate cellular environments have been crafted by hand (Gubins et al., 2021; Gubins et al., 2020; Xu et al., 2011). An algorithm targeting automated generation of cytoplasmic molecular arrangements at tunable crowding levels has been developed (Pei et al., 2016) but the approach is not suited to simulate membrane embedded molecular distributions.

Here, we introduce *MEPSi*, the *M*embrane *E*mbedded *P*roteins *Si*mulator. This tool was developed to allow quick and convenient simulation of membrane protein configurations embedded in membranes of tunable curvature. This model generation module is coupled with a tilt-series and tomogram generation module that is designed to allow rapid evaluation of imaging and data collection parameters either in combination or individually. While *MEPSi* can be used to generate charge densities for validating molecular recognition and sub-tomogram averaging and other approaches, its main utility lies in quick and exhaustive evaluation of imaging parameters in terms of interpretability and expected upper bunds for quality of individual raw tomograms.

## METHODS

### *MEPSi* workflow

*MEPSi* has been incorporated into *pyCoAn* (github.com/pyCoAn/distro), an extended python version of the CoAn package (Volkmann and Hanein, 1999). The *MEPSi* workflow is primarily accomplished with three commands within *pyCoAn*:

1. *sim_membrane_prots*. This command reads a coordinate list, generates the specified membrane geometry, places the proteins on the membrane, and creates a ground-truth volume representation of the resulting arrangement. The algorithm uses a three-dimensional Archimedean spiral to place the molecules initially at approximately equidistant points within the membrane. Parameters that can be adjusted include the membrane curvature, mean distance between molecules, the variance of random translations, whether a random rotation should be applied, the ratio at which the molecules will be randomly picked from the list, pixel size, resolution, and controls for the orientation and size of the output volume. Detailed help on all parameters is available within the *pyCoAn* software. The volume is generated using direct generation of the membrane density and Gaussian convolution of the atom positions. Optionally, solvent models can be generated and added to the density before tilt series simulation (*sim_add_solvent*).
2. *tilt_simulator*. This command reads a ground-truth volume, usually created by *sim_membrane_prots*, and generates an unperturbed basis tilt series with specified parameters. The individual tilt images are generated by rotating the volume around the Y axis (virtual tilt axis) and projecting the density along the Z axis (virtual electron beam direction). Tilt increment, tilt range, and pixel size can be adjusted. This step tends to be the most time-consuming within the workflow.
3. *tomo_simulator*. This command reads a basis tilt series, usually generated by *tilt_simulator*, and generates a simulated tomogram. It allows to adjust the thickness, tilt range and increment, pixel size, reconstruction algorithm, defocus, and whether noise should be applied and at what signal-to-noise ratio. Detailed help on all parameters is available within the *pyCoAn* software. The reconstruction is driven by *tomo3d* (Agulleiro and Fernandez, 2011) and includes options for the Simultaneous Iterative Reconstruction Technique (SIRT) and weighted back projection. This step is rapid and allows exploration of many parameters with the same base tilt series in a short amount of time.

Parameter modulations or other perturbations are applied to the basis tilt series. For example, different contrast transfer function (CTF) and/or noise models can be applied to the basis tilt series before selection (*sim_apply_ctf* and *sim_apply_noise* respectively). All *MEPSi* commands are completely modular and can be combined with other approaches. For example, a volume created by *sim_membrane_prots* could be used as input to a different, tilt-image generator and the resulting tilt series can then be fed back into *tomo_simulator* or hand-crafted volumes can be input to *tilt_simulator* followed by *tomo_simulator*.

*MEPSi* is agnostic to the source of the coordinate files (e.g., data base, prediction, hand-crafted, MD simulations, etc.) but there are two requirements for all coordinate files that need to be met: the center of the transmembrane domain needs to be at the origin of the coordinate system and the extracellular domain needs to point down the Z axis. *pyCoAn* provides commands to adjust the geometry of arbitrary coordinate files.

### Proteins targeted in this study

The four full-length αIIbβ3 integrin conformers were obtained previously and described in detail (Xu et al., 2016). Briefly, models were obtained by combining normal-mode-based and statistics-based docking to fit the different domains extracted from crystal structures into cryo-EM reconstructions. The coordinates of the SARS-CoV-2 spike conformers were obtained from the PDB with accession numbers 6VSB (pre-fusion conformation) and 6M3W (post-fusion conformation).

### Image quality assessment methods

Three image quality assessment methods were set up in this pipeline and implemented within *pyCoAn*: Mean Square Error (MSE), Peak Signal to Noise Ratio (PSNR) and Mean Structural Similarity Index Measure (MSSIM). Because none of these measures is intensity-scale invariant, each simulated tomogram was normalized to a mean value of zero and a root-mean-square deviation of one to facilitate meaningful comparison.

### Visualization

Experimental and simulated noise images, as well as slices and orthogonal slice representations of the simulated tomograms were visualized with IMOD (Kremer et al., 1996). Surface representations of the densities and atomic models were visualized with UCSF Chimera (Pettersen et al., 2004).

## RESULTS AND DISCUSSION

We developed *MEPSi* as a fast and easy-to-use simulation approach with the primary aim to determine and optimize cryo-ET data collection parameters (for example tilt range, tilt increment, defocus) for transmembrane proteins in their native membrane environment, The pipeline developed here includes importing of protein coordinates, rapid charge density creation, tilt series generation, and tomogram simulation (**Figure 1)**. In a few steps and in a short time, several tomograms with different characteristics can be simulated. The most time-consuming step of the pipeline is the generation of projection images for the tilt series from a charge density. To speed up the subsequent processes, a perturbation-free “basis tilt series” is generated from a particular charge density with small pixel size (default: 2 Å), small tilt increment (default: 1°), and large tilt range (default: ±90°). The simulated imaging and data collection parameters to be explored (along with perturbations such as noise or miss-alignment) are then applied to the images in the basis tilt series before or during tomogram calculation.

**Figure 1.**
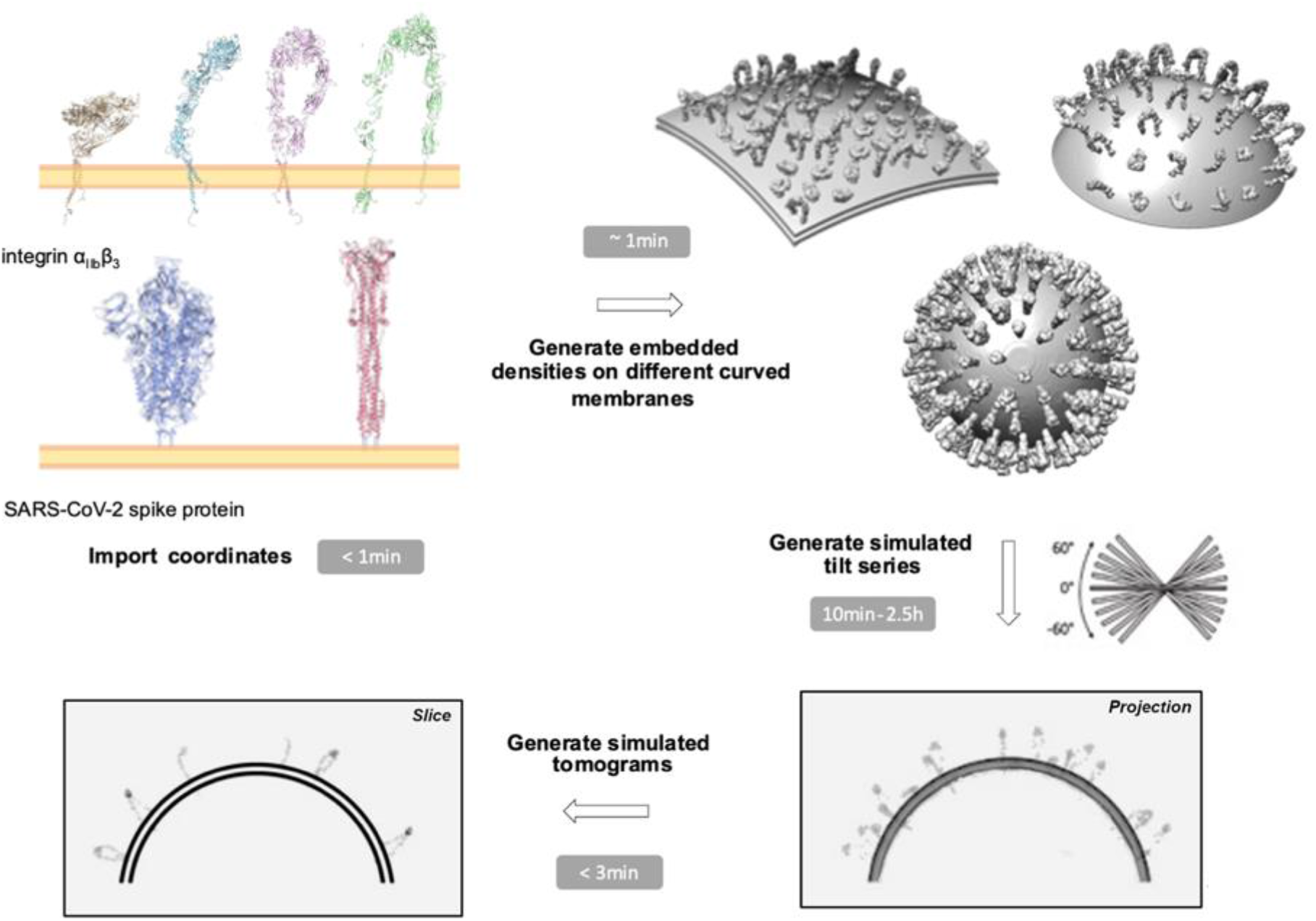
MEPSi (Membrane Embedded Proteins Simulation) workflow. The first step consists of importing and converting atomic coordinates, which takes a few seconds. The second step consists of generating the molecular arrangement from the imported coordinates, adding the membrane contribution with a defined curvature and, optionally, adding a solvent model with or without structural noise. This step typically takes few minutes, depending on the size of the volume. The next step consists of generating a “basis tilt series” that can be used to generate sub-sampling for the spatial parameters tilt range and increment as well as pixel size. It needs to be created with the finest tilt increment and pixel size relevant to the simulation aim and with the largest tilt range needed. This step is the most time-consuming, typically ranging between 10 mins and 2.5 hrs, but only needs to be done once per simulated volume. Once the basis tilt series is generated, the effects of parameters on the tomogram quality can be explored quickly, either together, or separately. Currently implemented parameters to explore at this level include tilt range, tilt-increment, pixel size, signal-to-noise ratio, tilt-series alignment accuracy, acceleration voltage, defocus, amplitude contrast, spherical aberration, and envelope function fall-off. Impact of packing, membrane curvature, specimen orientation in respect to the beam, solvent density and structural noise can also be easily explored but require recalculation of the basis tilt series.

### Charge density modeling

Under certain conditions, the image formation in the electron microscope can be considered as a linear image model (Thon, 1971). For this to hold, the specimen needs to be very thin and composed of light atoms. This is referred to as the weak phase object approximation where the effect of the specimen on the incoming electron wave is modeled as a spatially varying phase shift in the electron wave as it passes through the specimen, modifying the exit wave accordingly. The projection approximation assumes that the exit wave can be computed as the projected interaction potential of the whole specimen (Kirkland, 2010). Both approximations greatly simplify the computational complexity of simulating images but cannot account for effects like the curvature of the Ewald sphere (DeRosier, 2000) or multiple scattering events and are thus not suitable for very high resolution or thicker specimen.

Single isolated atoms with low to medium atomic number satisfy the conditions for both the weak-phase object and the projection approximation and can be treated accordingly. Electron scattering by an atom starts with a plane wave incident on the atom which gives rise to an outgoing plane wave plus an outgoing spherical wave, assuming spherical symmetry of the atom (an assumption that is technically broken in molecules). The amplitude of this exit wave is referred to as the electronic scattering factor. Following the first Born approximation, the Fourier transform of the scattering factor can be used to calculate the radial atomic potential (Kirkland, 2010). Those scattering factors can be parametrized by a weighted sum of five Gaussians (Peng et al., 1996). The charge density of a thin enough biological specimen can then be approximated using the isolated atom superposition approximation, which neglects the minor effects of the bond contributions to the interaction potential (Vulović et al., 2013).

All these approximations speed up calculations significantly if compared to more accurate treatments and are the standard implementation for conversion of atom coordinates into charge densities in most cryo-EM software packages. However, for large areas of crowded molecules such as those in cellular environments, even these approximate calculations tend to become a bottleneck for simulations. Although the number of electrons increases with the atomic number, the actual size of the atoms is surprisingly similar. The increased charge of a nucleus with higher atomic number attracts the electrons more strongly to the nucleus, keeping the atomic diameter relatively constant (~1 Å). Thus, directly approximating the radial charge distribution of an arbitrary atom as a point scatterer convoluted with a single Gaussian should hold at low resolution.

In *MEPSi* we approximate all atoms as point scatterers with equal weights and convolve the resulting point-scatterer distribution with a single Gaussian that incorporates the diameter of the atoms, fall-off for limited resolution, and, optionally, all sources of uncertainties. This strategy leads to a major improvement in computational load. To quantify the penalty in accuracy, we compared the results of the full-fletched five-Gaussian approach with our own approach. We find that the difference in the resulting charge distributions is negligible down to pixel sizes of at least 1.4 Å (the Fourier shell correlation stays above 0.5 up to 2.8 Å and above 0.143 up to 2.6 Å), especially in the presence of noise and imaging artifact such as the missing wedge. Once calculated, the resulting charge density can then be combined with continuum models for membranes and bulk solvent. Pixel sizes in cellular tomography are most often chosen between 3.5 and 10 Å to maximize the field of view unless high-resolution sub-tomogram averaging is targeted. New technology potentially allows the recording of large fields of view (Peck et al., 2022) with smaller pixel sizes but this technology is not widely used yet. Here, we chose a pixel size of 2 Å for the analysis as a compromise between current practice and future prospects.

### Molecular packing

One significant issue is how to get the packing right in crowded environments. In three dimensions, it can be resolved in several different ways including simulated annealing and molecular dynamics approaches (Pei et al., 2016). For membrane-bound macromolecules, the problem reduces to packing consideration on a two-dimensional manifold. Our approach to this issue is to first place the macromolecules in a predefined near-equidistant pattern using a three-dimensional Archimedean spiral approach and then apply random translations on the surface within a bounding box defined by the equidistance used and the maximum XY radius of the respective macromolecule. This approach ensures that there is no overlap between the macromolecules on the surface.

### Solvent modeling

The contrast in experimental images is generally much less, typically by about a factor of three, than in simulated images using multi-slice approaches (Hÿtch and Stobbs, 1994). This issue has been addressed recently by using sophisticated frozen plasmon forward modeling (Himes and Grigorieff, 2021) but the improvement comes at a considerable computational cost. Instead, we use a continuum solvent model with an adjustable contrast tuning parameter. As an added benefit, this strategy allows modeling the effect of denser buffers and or graphene films that generate lower contrast than the pure water that is generally used in other biological specimen simulators. Optionally, random bumpiness can be introduced into the solvent model to emulate structural noise, which has been shown to be an important factor in signal-to-nose ratio considerations (Baxter et al., 2009). To combine the solvent, membrane, and macromolecular charge density models, we use a three-dimensional version of Laplacian pyramid blending (Burt and Adelson, 1987) to account for displacements of one object from another and to mitigate edge effects. This type of blending also effectively emulates the existence of a hydration layer around the molecular objects (Shang and Sigworth, 2012).

### Contrast transfer modeling

The effect of the microscope optics on the simulated image can be modeled with the contrast transfer function (Thon, 1971). One major determinant of the CTF is the focus value at the scattering event. One of the reasons why the projection and weak-phase object approximations break down for thicker samples is the fact that the focus value changes while the electrons traverse the specimen. This effect is even more relevant for tilted specimen, where the focus difference can reach several microns in magnitude at high tilt angles. The maximum resolution *R* achievable from a specimen of thickness *t* while employing the projection approximation can be expressed as 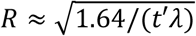 (Philippsen et al., 2007) where *λ* is the wavelength of the electrons and *t’* = *t/cos(α)* is the tilt-corrected thickness at tilt angle *α*. For an acceleration voltage of 300 kV and a thickness of 250 nm, at the higher end of thickness for cellular lamellae obtained by recent cryogenic focused ion beam milling approaches (Tacke et al., 2021), the reliable information is restricted to a resolution of 5.5 Å at zero tilt and 7.8 Å at 60° tilt. The value for the high tilt is above Nyquist limit for any pixel size larger than 3.9 Å and employing the projection approximation for pixel sizes down to 2 Å should not be a major factor considering the low signal-to-noise ratio in tomograms and the general magnitude of the imaging artifacts.

We exploit this fact by treating the simulated specimen as an infinitely thin slice so only focus changes caused by tilting need to be considered. We implement this procedure as a two-dimensional version of a multi-slice approach, where the projected tilted specimen images are subjected to a CTF model in strips parallel to the tilt axis with the defocus modulated according to the position of the strip center in the three-dimensional tilted specimen. The width of the strips is tunable using the desired defocus step size and strips are blended into a final image using Laplacian pyramids. This approach accelerates the calculations significantly if compared to the traditional multi-slice approach.

### Noise modeling

In many simulation applications targeting cryo-ET, the noise component is modeled as white additive Gaussian, which has been demonstrated to be inadequate in accounting for common imaging artifacts during image processing (Scheres et al., 2007). Here, we model noise as a mixture of Gaussian and Laplacian noise with tunable mixing and noise-correlation parameters. Tuning these parameters allows to closely match the simulated noise with experimentally observed noise (**Figure 2**). This noise is then applied to the simulated tilt images at defined signal-to-noise ratios before tomogram generation.

**Figure 2:**
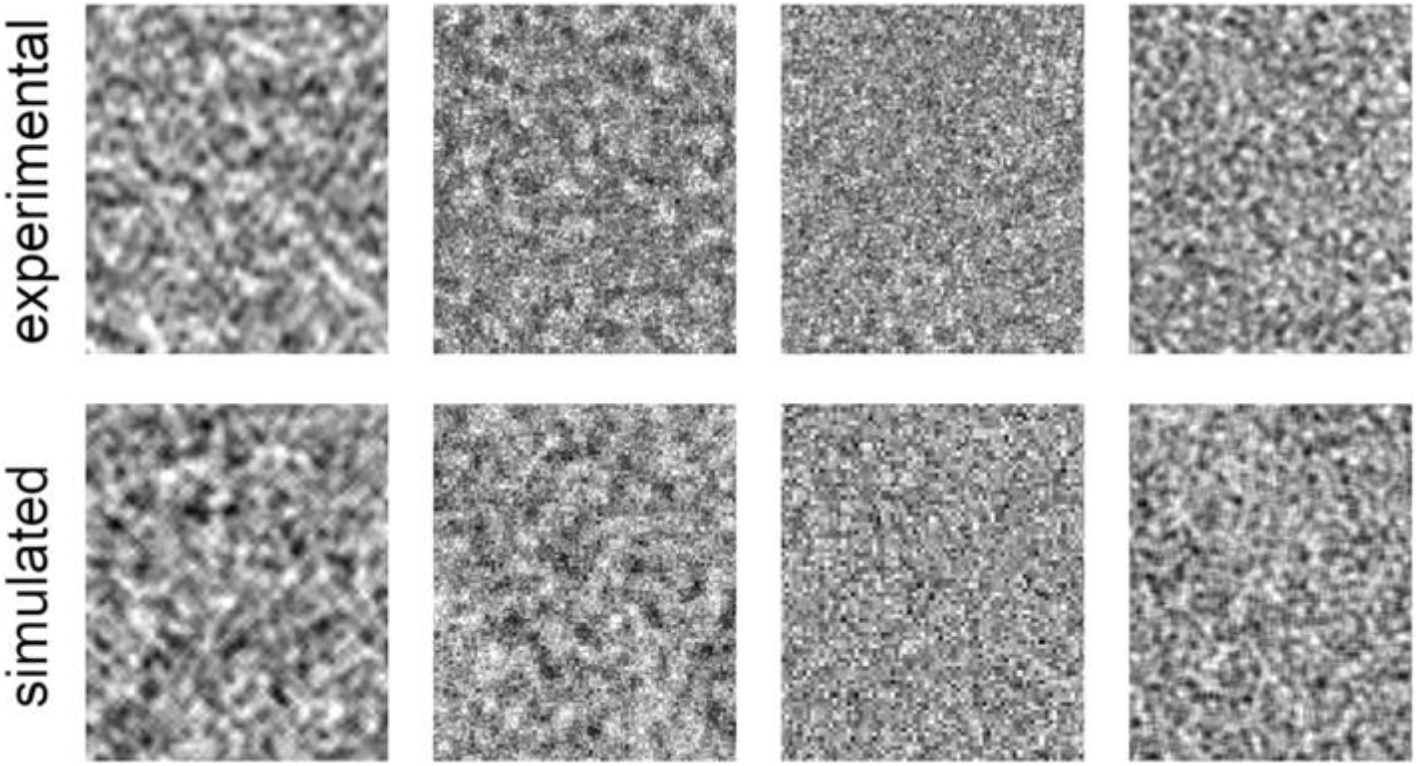
Noise modeling using mixtures between Gaussian and Laplacian noise with tunable mixing and noise-correlation parameters. The top row shows areas of experimental electron micrographs from different experiments with different imaging parameters in regions devoid of structures (which should thus only show noise). The bottom row shows the results of the noise simulations with parameters adjusted to match the experimental noise.

### Processing speed

The generation of the basis tilt series can take up to 2.5 hours with default parameters for a 2000 x 2000 x 500 tomogram on a standard workstation. This time can be reduced to less than 10 minutes while still giving a good visual indication for the quality of the resulting tomograms by reducing the size of the tomogram and/or the pixel size before calculating the basis tilt series. The latter time frame makes *MEPSi* amenable to real-time evaluation during data collection. In either case, once the basis tilt series is generated, tomograms with different parameter settings can be generated in a matter of minutes. The algorithms are optimized for multi-CPU usage and do not depend on the presence of graphic cards.

### Model systems

One of the main features of *MEPSi* is that it allows easy generation of different charge density geometries and compositions (**Figure 3).** Membrane curvature (**Figure 3A**), the spatial distribution and concentration of proteins on the surface (**Figure 3B**), and ratios within the conformational landscape (**Figure 3C**) can all be easily adjusted. To demonstrate *MEPSi’s* utility we employed *MEPSi* to characterize the individual influences of the parameters *tilt range* and *tilt increment* for two different proteins, the αIIbβ3 integrin receptor, and the SARS-CoV-2 spike protein. For reconstituted systems of αIIbβ3 integrin there is information available from X-ray crystallography and Nuclear Magnetic Resonance spectroscopy (Campbell and Humphries, 2011) as well as from cryo-EM (Xu et al., 2016; Ye et al., 2008). Only little *in-situ* structural information has been obtained so far (Sorrentino et al., 2021). For αIIbβ3 integrin, it is particularly important to be able to map individual conformations and locations on the plasma membrane because integrin function is governed by conformational equilibria (Hanein and Volkmann, 2018) and spatial clustering (Welf et al., 2012). Whether it is possible to extract this information from cellular tomograms directly and what the optimal imaging parameters are to achieve this goal is not known. For SARS-CoV-2 spike protein in its native virus context, near-atomic information after sub-tomogram averaging is available (Turoňová et al., 2020), demonstrating that it is possible to extract high-resolution information from *in-situ* viruses if sample and imaging conditions are right. However, a systematic analysis of what imaging parameters would lead to the minimum investment in microscope time and image analysis effort is not available. *MEPSi* opens the avenue for optimizing parameters in both cases.

**Figure 3.**
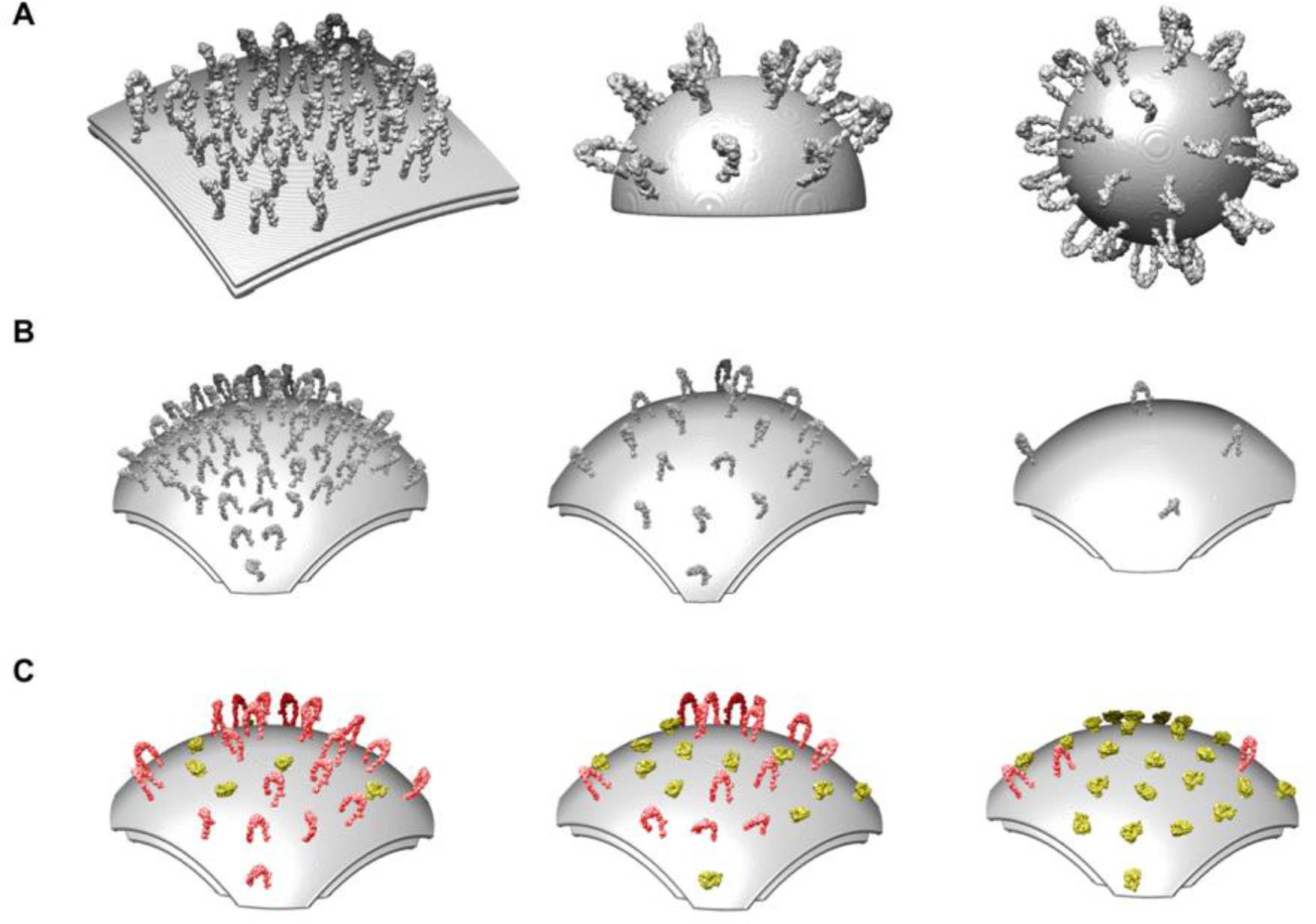
Tunning membrane curvature, protein densities and composition. **A)** Curvature variation. From left to right: underlying radius 3200, 1200 and 400 Å. **B)** Tunning of protein density on the membrane. From left to right: mean distance between particles 170, 300, and 550 Å. **C)** Variation of protein ratios considering two different conformations of integrin, (open conformation in red and closed conformation in yellow). From left to right: ratios of 75/25, 50/50, and 25/75.

As a proof of concept, we simulated different charge densities with two membrane geometries (flat and curved) for *in-situ* full length αIIbβ3 integrin arrangements. The ‘curved’ membrane geometry corresponds to integrins at the tip of filopodia (Sorrentino et al., 2021) and the flat geometry corresponds to the arrangement on extended flat membrane regions at sites like focal adhesions. For SARS-CoV-2, we only generated one membrane configuration that corresponds to a spherical shape with the known virus diameter of around 90 nm (Klein et al., 2020). For the αIIbβ3 integrin simulations, we drew from the four conformers seen in nanodiscs using the percentages from the conformational equilibrium of the activated state (Xu et al., 2016). Because the effect of parameter changes will be different for different orientations, we simulated tomograms with three different orthogonal orientations for each of the curved or flat configurations. The basis tilt series were calculated at a tilt increment of 0.25° and ±90° tilt range. The direction of the tilt axis coincides with the Y axis and the direction of the electron beam (i.e., the projection direction) coincides with the Z axis. Because the membrane of SARS-CoV-2 viruses are spherical, there was no need to generate tomograms of more than one orientation. We used a ratio to 3:1 for post-fusion versus pre-fusion states of the SARS-CoV-2 spike protein.

For each simulated tomogram, we varied the tilt increment between 1° and 5° and the tilt range between ±90° (no missing wedge) and ±45°. Effects of these parameters are visually summarized for the flat αIIbβ3 integrin configuration with the membrane oriented parallel to the Z axis in **Figure 4**. In this configuration, the contrast of the membrane in the simulated tomograms is significantly diminished once a missing wedge is introduced. The membrane practically disappears at small tilt range (±45°). A tilt range of ±70° provides a clear improvement in membrane visibility but there are significant distortions in the location of the membrane (**Figure 4B)**. Interestingly, modifying the tilt increment does impart a significant effect even without introducing a missing wedge (**Figure 4C**). For a tilt increment of 5°, the membrane is severely distorted in the view along the Z axis and there are significant streaks in the other two views. The tomogram calculated from 3° tilt increments is less severely affected but the sampling artifacts are still of the same magnitude as the signal. A 2° tilt increment appears to be sufficient to suppress sampling artifacts to a tolerable level. Smaller tilt increments are impractical because of the electron dose will become so low that alignment can become unstable. Taken together, a ±70° tilt range with a tilt increment of 2° is the best choice for this geometry.

**Figure 4.**
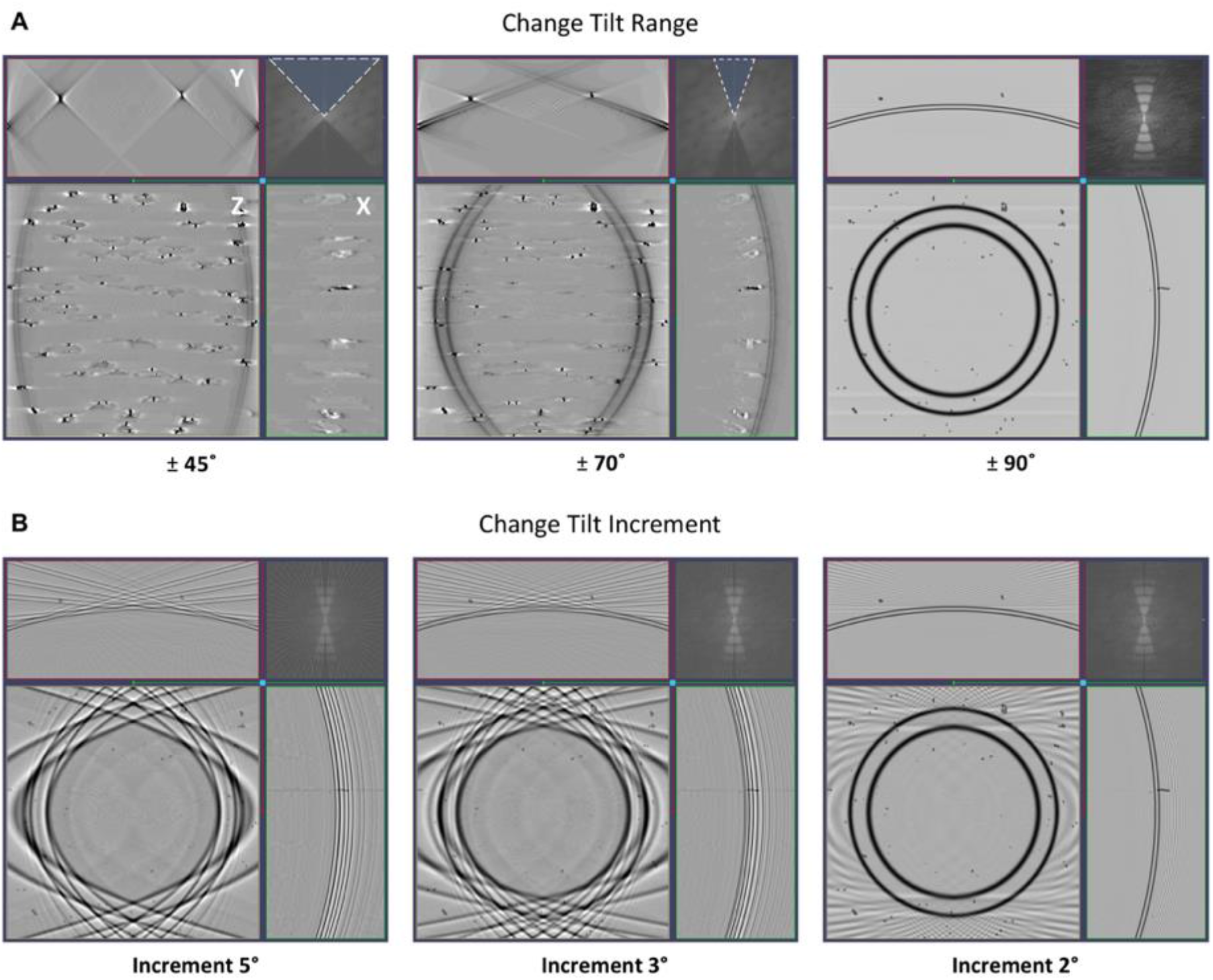
Simulated tomograms. **A)** Orthogonal slices representations of simulated tomograms with different tilt ranges (tilt increment 0.25°) and the membrane oriented perpendicular to the electron beam. Z denotes the view along the direction of the electron beam; Y denotes the view along the direction of the tilt axis; X. The inset in the top right corner of every orthogonal slice representation shows the Fourier transform of the central slice along the tilt axis. The missing wedge is highlighted in blue with a dashed white border. **B)** Orthogonal slice representations of simulated tomograms with different tilt increments (tilt range ±90°). Axis and insets as in A.

### Quantification

An essential component of evaluating different parameter choices is quantification of the differences in quality. A good quality metric will not only allow to track how the choice of a particular parameter affects the tomogram quality, it will also allow to quantitatively compare the effect of different data collection or imaging parameters. We have implemented three reference-based image comparison methods that are popular in computer vision: (1) the Mean-Square Error (MSE), which measures the average squared difference between the simulated values and the actual values (ground truth); (2) the Peak Signal-to-Noise Ratio (PSNR), which measures the ratio between the maximum possible value of a signal and the strength of the distortions; and (3) the Mean Structural Similarity, (MSSIM), which attempts to quantify the image degradation as the change of perception in structural information and is thought to better match human visual quality perception than the other two metrics (Zhou, 2004). All three metrics reveal clear and mostly similar trends as a function of varying parameters (**Figure 5)**. Interestingly, for many membrane geometry/orientation combinations the tilt increment has as much or more influence on the metrics as the tilt range. The one exception is the flat membrane geometry when oriented perpendicular to the electron beam (flat Z). Here, the MSE is almost an order of magnitude higher at ±45° tilt range than the MSE of the other geometry/orientation combinations. The tilt increment dependence is comparable to the others. The reason for the high MSE is that the membrane all but disappears from the reconstructions when the missing wedge grows, leading to a large pixel-wise difference in the membrane regions. Because both the PSNR and the MSSIM are considering localized signal strength, they are not picking up as much on the absence of the membrane, indicating that these metrics are more focused on the appearance of the particles in the reconstruction.

**Figure 5.**
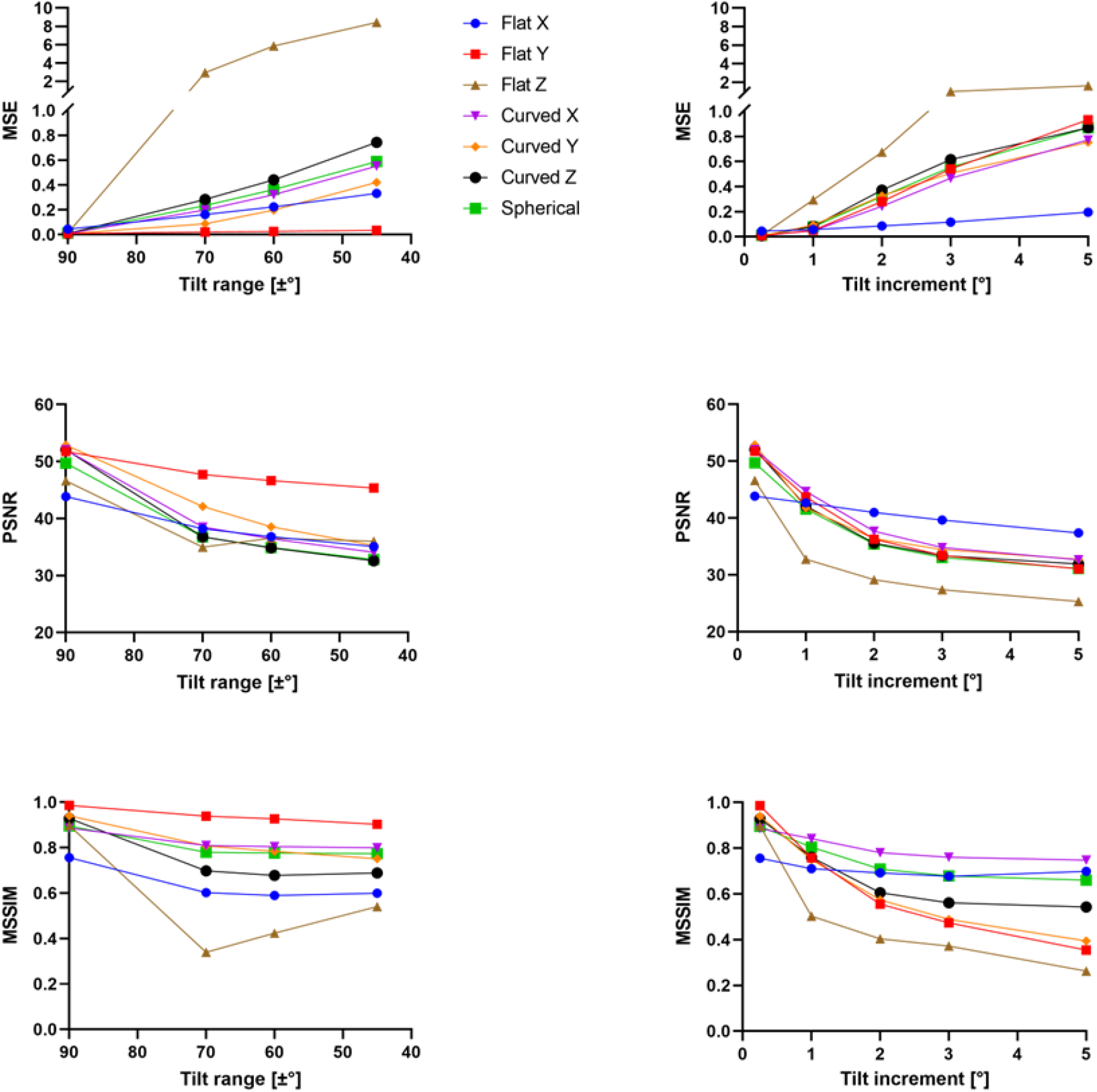
Assessing image quality of tilt range and tilt increment in simulated tomograms with different membrane geometries. MSE, PSNR and MSSIM as a function of varying parameters. For the integrin-based densities two membrane geometries (Flat and Curved) and three orthogonal orientations were evaluated whereas for the virus-like structure, which has a spherically symmetric membrane, only one orientation was evaluated (Spherical). X denotes that the membrane is oriented perpendicular to the tilt axis and parallel to the electron beam, Y denotes that the membrane is oriented parallel to both the tilt axis and the electron beam; Z denotes that the membrane is oriented perpendicular to the electron beam.

## CONCLUSIONS

In cryo-ET one of the most difficult problems is the interpretation of the data after reconstruction. In addition to the high noise levels and low contrast, the imaging system and the chosen imaging parameters significantly distort the imaged structures. *MEPSi* can help to visualize the extent of the expected distortions and aid interpretation of experimental tomograms. In particular, the orientation of the membrane in respect to the tilt axis and the electron beam can have a major influence on the interpretability of the reconstruction (**Figure 6**). If the membrane is oriented perpendicular to the tilt axis and parallel to the electron beam (i.e. oriented along the X axis), the overall distortions for a typical experimental setting (tilt range ±60°, tilt increment 3°) are significantly less disturbing than the other two membrane orientations. Clearly the worst scenario is when the membrane is orientated perpendicular to the electron beam. In this case, the membrane is not visible in either of the orthogonal slices.

**Figure 6.**
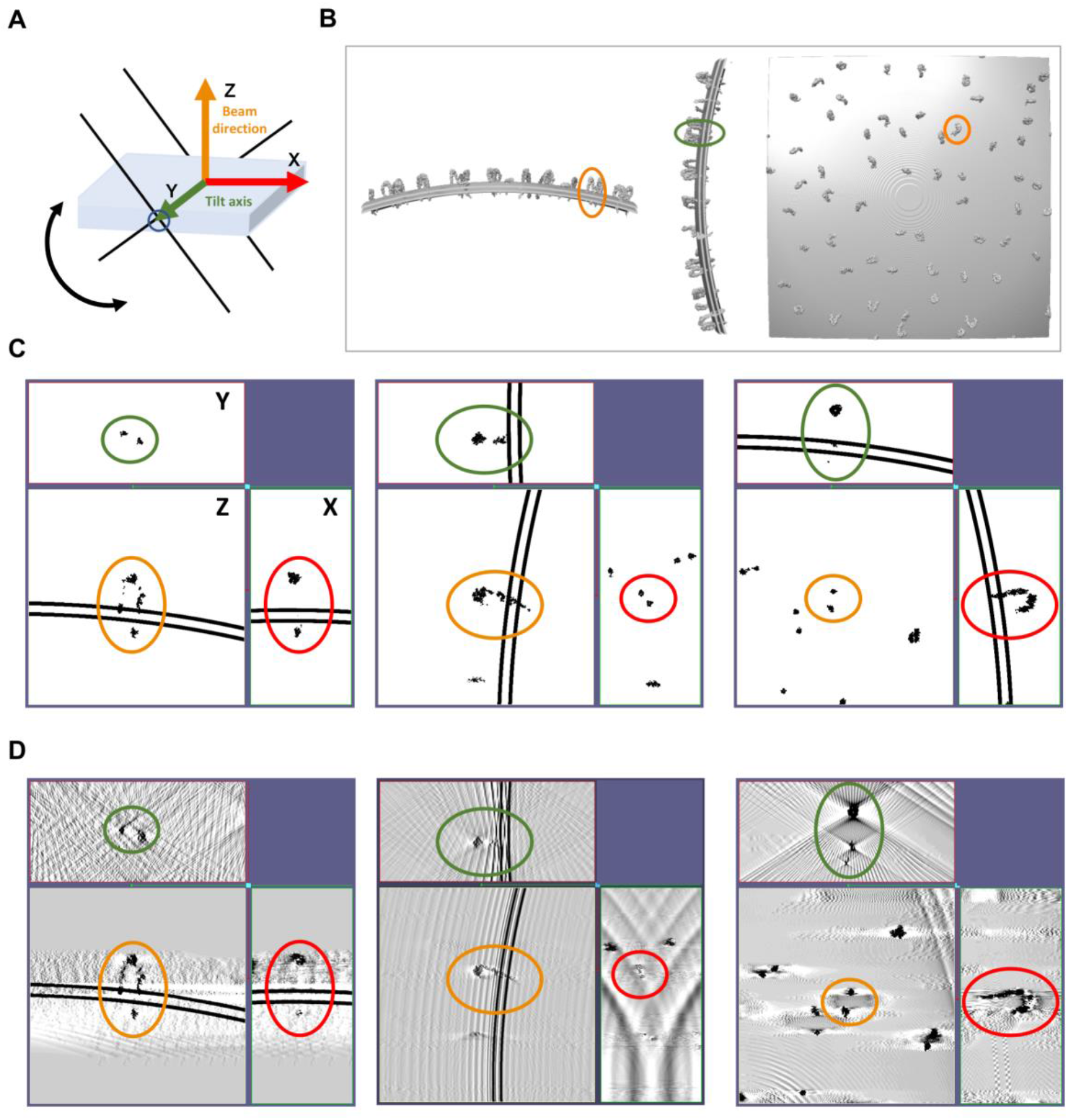
Influence of orientation on protein direct visualization. **A**. Schematic representation of the simulated geometry. **B.** Surface representations of charge density of integrins embedded in a flat membrane. Three orthogonal orientations were generated. All views along the Z axis (beam direction). The same integrin molecule is marked in all views. **C,** Orthogonal slice representations of the ground truth densities corresponding to the orientations shown in B. The same molecule as in B is circled. The view with the red circle corresponds to the view along the X axis, the one with the green circle to the view along the Y axis and the one with the orange circle to the view along the Z axis. **D.** Orthogonal slice representations through a simulated tomogram (tilt range ±60° and tilt increment 3°). The same molecule as in B is circled, color scheme as in B.

We demonstrated the utility of *MEPSi* for evaluating and comparing the effects of modifying data collection parameters for different membrane geometries. *MEPSi* is currently targeting membrane embedded molecules and is a versatile tool to explore the imaging and data collection parameter space as well as perturbations typical for cryo-ET such as high noise levels, structural noise, and misalignment of tilt images. Possible extensions of the method include adding capacity to simulate cytosolic distributions of macromolecules and/or filaments alongside the membrane embed molecules. In terms of additional parameters to explore, beam damage models, electron dose, and models for beam-induced distortions would be attractive. Another attractive option is the exploration of alternative reconstruction algorithms in our framework, Because of the modular design of *MEPSi*, it can also be adapted to use more accurate (albeit more time-consuming) forward models to validate and explore molecular recognition approaches or high-resolution sub-tomogram classification and averaging methodology.

## ABBREVIATIONS

Cryo-EM: Cryogenic Electron Microscopy
Cryo-ET: Cryogenic Electron Tomography
CTF: Contrast Transfer Function
MEPSi: Membrane Embedded Protein Simulator
MSE: Mean-Square Error
MSSIM: Mean Structural Similarity Index Method
PSNR: Peak Signal-to-Noise Ratio

## ACKNOWLEDGMENTS

The authors acknowledge the use of the Titan Krios, Tecnai Spirit T12 and auxiliary equipment at the cryo-EM unit of the Sanford Burnham Prebys Medical Discovery Institute. We thank Drs. Cécile Sauvanet and Fernando Vilela for valuable input to the manuscript and feedback during the development phase of the project.

